# Predicting Protein-encoding Gene Content in *Escherichia coli* Genomes

**DOI:** 10.1101/2023.01.17.524402

**Authors:** Marcus Nguyen, Zachary Elmore, Clay Ihle, Francesco S. Moen, Adam D. Slater, Benjamin N. Turner, Bruce Parrello, Aaron A. Best, James J. Davis

## Abstract

In this study, we built machine learning classifiers for predicting the presence or absence of the variable genes occurring in 10-90% of all publicly available high-quality *Escherichia coli* genomes. The BV-BRC genus-specific protein families were used to define orthologs across the set of genomes, and a single binary classifier was built for predicting the presence or absence of each family in each genome. Each model was built using the nucleotide k-mers from a set of 100 conserved genes as features. The resulting set of 3,259 XGBoost classifiers had a per-genome average macro F1 score of 0.944 [0.943-0.945, 95% CI]. We show that the F1 scores are stable across MLSTs, and that the trend can be recapitulated through sampling with a smaller number of core genes or diverse input genomes. Surprisingly, the presence or absence of poorly annotated proteins, including “hypothetical proteins”, were easily predicted (F1 = 0.902 [0.898-0.906, 95% CI]). Models for proteins with horizontal gene transfer-related functions, including transposition- (F1 = 0.895 [0.882-0.907, 95% CI]), phage- (F1 = 0.872 [0.868-0.876, 95% CI]), and plasmid-related (F1 = 0.824 [0.814-0.834, 95% CI]) functions had slightly lower F1 scores, but were still accurate. Finally, we applied the models to a holdout set of 419 diverse *E. coli* genomes that were isolated from freshwater environmental sources and observed an average per-genome F1 score of 0.880 [0.876-0.883, 95% CI], demonstrating the extensibility of the models. Overall, this study provides a framework for predicting variable gene content using a limited amount of input sequence data.

**Importance:** Having the ability to predict the protein-encoding gene content of a genome is important for a variety of bioinformatic tasks, including assessing genome quality, binning genomes from shotgun metagenomic assemblies, and assessing risk due to the presence of antimicrobial resistance (AMR) and other virulence genes. In this study, we built a series of binary classifiers for predicting the presence or absence of variable genes occurring in 10-90% of all publicly available *E. coli* genomes. Overall, the results show that a large portion of the *E. coli* variable gene content can be predicted with high accuracy, including genes with functions relating to horizontal gene transfer.

## Introduction

Having the ability to predict the protein-encoding gene content of a genome is important for a multitude of bioinformatic tasks. These include assessing the metabolic capabilities of a genome or community organisms sampled in a metagenome, estimating genome quality and completeness, and understanding the potential for a genome to encode antimicrobial resistance (AMR) and other virulence factors. Over the years, a variety of approaches for predicting protein-encoding gene content and their associated functions have been devised, and they vary based on the study design and the experimental needs. Some approaches are as straight forward as searching for conserved genes, while others use more complex algorithmic and artificial intelligence-based approaches to predict the presence or absence of key genes.

Perhaps the largest body of work predicting gene content has come from the field of metagenomics. Metagenomic studies routinely perform amplicon sequencing of the 16S rRNA gene in order to determine the microbial diversity of an environment. However, since 16S sequencing does not provide information about the gene content of the sample, many methods have been developed to infer gene content and functions from the 16S sequence. These include popular tools such as, PICRUSt (1), Piphillin (2), Tax4Fun (3), PAPRICA (4), PanFP (5), and MicFunPred (6). Although the algorithmic steps vary, in essence, these tools utilize the relationship between phylogeny and gene content and make their predictions based on a set of closely related reference genomes (7). These methods are useful for inferring the metabolic capabilities of the constituents of a sample, especially when deep shotgun sequencing is unavailable. However, they come with understandable limitations due to the limited information encoded in 16S amplicon sequences (particularly single variable regions), the size and scope of the reference databases, and the ability of horizontal gene transfer to disrupt gene content between close relatives.

Another important bioinformatic application, the prediction of genome completeness—i.e., knowing that all genes are present in a given genome—is crucial for assessing genome quality and the reliability of downstream comparative analyses (8). This is important when working with genomes that are assembled from short reads, genomes from single cell sequencing data, and metagenome assembled genomes (MAGs). Many studies have established measures of genome completeness by searching for sets of universal or lineage-specific genes from close relatives including CheckM (9), BUSCO (10), CEGMA (11), mOTU (12), Anvi’o (13), and CheckV (14). Parrello and colleagues recently extended this concept by building a tool that predicts genome completeness for bacterial and archaeal genomes from the protein annotations in the PATRIC database (15, 16). Using a set of approximately 2,000 well annotated “roles,” which are the individual atomic functions of a protein in the SEED annotation schema (17), they built a set of machine learning classifiers that predicted the presence or absence of each role based on the presence or absence of the other roles in the set. This enabled them to both quantify the completeness of the genome and provide an estimate for the expected number of occurrences of each role per genome. In most cases, when genome completeness scores deviate from expectation, the genome is reliably incomplete or contaminated with sequences from another organism. Similarly, recent updates to the CheckM algorithm, CheckM2, incorporate the use of machine learning models, which include the KEGG protein annotations as part of the feature vector (18, 19). Another recent tool called MetaPredict uses a set of classifiers to predict the presence or absence of KEGG modules in a MAG, based on the presence or absence of the existing annotations in the MAG (20). Although all of these genome completeness methods have proven to be useful for assessing genome completeness, a potential downside is their reliance on collections of well characterized genes, which may not fully capture patterns in the variable strain-specific gene content across a species.

AMR genes and other virulence factors are often among the set of strain specific genes that vary between the members of a species. Many of these genes are found on mobile genetic elements, so their occurrences sometimes do not match the phylogeny of a given taxon. Many bioinformatic tools have been developed to search for AMR and virulence genes within a genome or metagenome using both sequence similarity (21–28) and machine learning techniques (29, 30). Since shotgun metagenomic studies sample multiple genomes, and their assemblies are often incomplete, methods that identify AMR and virulence genes are not always able to identify the source genome for a given AMR gene. To this end, some studies have attempted to predict the source genomes for the AMR genes in a sample using either statistical (31) or machine learning methods (32, 33).

Many studies have also been designed to predict AMR phenotypes from genome sequences by training machine learning models using the genomes and laboratory-derived antimicrobial susceptibility test data (34). Importantly, several of these studies have demonstrated that AMR phenotypes can be predicted using the phylogeny of the strains, either by learning the tree structure, mapping phenotypes from close relatives, or building machine learning models from conserved parts of the genome (35–38). This has been demonstrated even in cases where the AMR phenotype is the result of a horizontal gene acquisition and is presumably due to the machine learning models learning non-linear relationships in the input data. Although there is a clear link between phenotype and genotype, to date we still lack well-developed tools for predicting whether an AMR or virulence gene should or should not be present in a genome given a set of existing sequences from a contig or MAG.

*E. coli* is the most widely studied bacterial species, and there are currently well over 30,000 sequenced *E. coli* genomes in the public domain. All of the genes of the species can be thought of as a pan-genome consisting of a conserved set of core genes that are held in common among all members of the species, plus tens of thousands of accessory genes that are often strain-specific and vary in their frequency of occurrence (39–43). These variable genes encode a variety of known and unknown functions including many AMR and virulence genes. In this study, as a proof of concept, we wanted to see the extent to which it is possible to predict the presence or absence of variable genes in *E. coli* using the nucleotide sequences of a core set of universal genes to make the predictions.

## Materials and Methods

### Genomes and datasets

A high-quality, diverse set of publicly available *Escherichia coli* genomes was selected for building the models. All *E. coli* genomes were downloaded from the Bacterial and Viral Bioinformatics Resource Center (BV-BRC) FTP site (ftp.bvbrc.org) on March 28, 2022. The BV-BRC is a large bioinformatics resource center that maintains the PAThosystems Resource Integration Center (PATRIC)(15, 44). Each bacterial genome in the BV-BRC has been uniformly annotated using the Rapid Annotation Using Subsystem Technology (RAST) pipeline (45), and the analysis includes computation of genome quality (16), protein encoding gene annotations (45), protein family assignments, etc. (46). All *E. coli* genomes lacking “Complete” or “WGS” designations (sourced from GenBank) (47), and those that were listed as being poor quality (16), were excluded from consideration. Any genome with less than half of the average number of genes per genome was also excluded. The genome set was further filtered to ensure that all of the core genes that were used for generating features for the models (described below) were present, and that each gene with a given function was within 50-200% of the median gene length. This resulted in a set of 34,527 *E. coli* genomes that were available for modeling.

In order to reduce the size of the set of genomes for computing efficiency, while maintaining genomic diversity, genomes were clustered based on nucleotide k-mer similarity. A set of 100 core genes, defined as those corresponding to the protein families that were most highly conserved across the entire set of *E. coli* genomes, was computed (**Table S1**). Nucleotide 7-mer counts were computed for the core genes of each genome using KMC 2.3.0 (48) and the genomes were clustered based on their 7-mer distances using the agglomerative clustering function in scikit-learn (version 0.20.3) (49) using the parameters: n_clusters=’4000’, affinity=’l1’, and linkage=’average’. From this, we selected a final set of 4,000 diverse *E. coli* genomes representing each cluster that was used for training and testing the models in this study (**Table S2**).

Since the goal of this study was to predict whether a protein-encoding gene should be present or absent within a given *E. coli* genome, we chose to use the PATRIC local protein families (PATtyFams) to describe this set (46). Genes encoding proteins that were members of the same protein family were considered to be orthologous. We computed the frequency of occurrence for each protein family across the entire starting set of 34,527 *E. coli* genomes and chose to model the protein families occurring in 10-90% of the genomes. This resulted in a total of 3,259 *E. coli* protein families that were modeled (**Table S3**). We chose not to model protein families occurring in less than 10% of the genomes to avoid class imbalance and to keep the number of models tractable.

### Model generation

The set of 100 nearly universal core genes (described above) was chosen for generating the k-mer based feature sets for the models (**Table S1**). The genes were found in each *E. coli* genome, and the nucleotide sequences were subdivided into canonical 7-mers using KMC version 2.3.0 (48). 7-mers were chosen because they train rapidly while retaining accuracy (**Table S4**). A matrix was created where the columns were the k-mers, the rows were the genomes, and each cell contained the counts of each k-mer. K-mers containing ambiguous nucleotides were not considered. A binary classifier was computed for each of the 3,259 protein families described above, where the labels were the presence or absence of the family in each genome.

Models were built using Extreme Gradient Boosting (XGBoost) version 0.81 (50) as described previously (36, 51). Unless otherwise stated, all models were evaluated using a 10-fold cross validation, where 80% of the data were used for training, 10% for testing, and 10% as a holdout set to monitor for overfitting in each fold. Model parameters were chosen based on tuning experiments for conserved gene models that were previously published (36). These included a maximum tree depth of 16 and a learning rate of 0.0625. Due to the high computing volume, unless otherwise stated, results are shown for the first five of ten folds.

### Environmental genomes

A holdout set of genomes from 419 environmental *E. coli* isolates was used to evaluate the models that were trained on the public genomes (**Table S5**). *E. coli* isolates were collected from freshwater samples in rivers, streams, and Lake Macatawa in the Macatawa Watershed (Holland, Michigan, USA) between 2012 and 2019 as part of year-round water quality monitoring efforts. EPA Method 1603 (52) was used to monitor *E. coli* levels in the watershed and served as the basis for strain collection. Isolated colonies displaying morphology consistent with *E. coli* on mTEC plates were streaked for isolation on nutrient agar plates to obtain pure cultures of putative *Escherichia* strains. Purified isolates were archived as glycerol stocks and stored at −80°C for downstream genome sequencing. All strains were screened via standard biochemical identification tests to ensure consistency with *E. coli* phenotypes prior to sequencing. Genomic DNA extraction was performed with the DNeasy® PowerLyzer® Microbial Kit (Qiagen). Sequencing library preparation was performed with the Nextera XT DNA Library Prep kit (Illumina). Library QC was performed with the Qubit™ dsDNA HS Assay Kit (Invitrogen) and an Agilent 2200 TapeStation system, using the High Sensitivity D5000 ScreenTape System (Agilent). Pooled libraries (24 per run) were sequenced on an Illumina MiSeq using the MiSeq Reagent Kit V2 (500 cycle, PE 2×250), according to manufacturer instructions. Genomes were assembled and annotated using the BV-BRC assembly and annotation services (44).

### Subset analyses

Several experiments were conducted to determine how models performed with less data. In order to evaluate model performances on a smaller number of genomes, clustering (as described above) was performed to generate sets that were 500, 1000, 2000, and 4000 genomes in size, and modeling was subsequently performed on these representative genome sets. To evaluate model performances using fewer conserved protein families, the top 25, 50, and 75 most conserved genes were selected from the original set of 100 conserved genes (**Table S1**) and models were trained on each respective set. Model performances were then recorded as described above.

### Genomic comparisons

Multi Locus Sequence Types (MLST) were computed for all genomes using the MLST tool version 2.21.0 developed by Torsten Seemann (https://github.com/tseemann/mlst), which uses the PubMLST database (53). The phylogenetic tree was computed based on a concatenated nucleotide sequence alignment of the genes corresponding to the five most conserved protein families in **Table S1**. Genes were aligned using MAFFT v7.130b (54). The alignment was curated by removing all inserts occurring in less than 5% of the genes, and poor quality sequences were removed by hand using the alignment editor JalView version 2.11.2.0(55). A tree was generated with FastTree version 2.1.7 using the generalized time reversable model for nucleotide sequences (56). Trees were rendered in iTOL (57). *Salmonella enterica* serovar Typhimurium LT2 was used as an outgroup for the tree (GenBank ID: AE006468.2).

### Data availability

Genomes for environmental isolates have been deposited at SRA under bioprojects PRJNA923802 and PRJNA918992 . Modeling software is available on github https://github.com/BV-BRC-dependencies/EColiVariableGeneModels.

## Results

### Predicting variable gene content

In order to predict the presence or absence of variable genes across the set of *E. coli* genomes, we first determined a set of genes that were amenable to modeling. To do this, we defined orthologous genes as those that belong to the same PATRIC local protein family (46) across the *E. coli* genomes in the BV-BRC database (44). The local protein families are restricted to each genus, and they are computed using the same set of signature amino acid k-mers that are used by the RAST annotation system to project protein functions (46). The protein family algorithm also considers all “hypothetical proteins”, placing them into families using either signature k-mers or sequence similarity with BLAST (58), thus enabling the tracking of the these poorly annotated genes. We chose to exclude highly conserved genes occurring in greater than 90% of the genomes and rare genes occurring in less than 10% of the genomes because of the difficulty in building balanced models to predict their presence or absence. This resulted in a final set of 3,259 variable genes occurring in 10-90% of the *E. coli* genomes that were modeled in this study (**Figure 1**). The set of genes encodes a diverse set of proteins with a variety of strain-specific functions (**Table S3**). Overall, 679 of the genes encode proteins with functions that exist in a curated SEED annotation subsystem (17). Some of the more common functions include components of secretion systems, fimbriae and flagella, toxins and antitoxins, and genes involved in transcriptional control. Many have annotations relating to horizontal gene transfer (e.g., phage, transposition, and plasmid conjugation related functions). Over 40% of the genes encode proteins with poorly annotated functions containing the terms, “hypothetical,” “uncharacterized,” “putative,” or “mobile element protein” (we note that in the SEED annotation schema the term “mobile element protein” is an outdated term that is more often synonymous with “hypothetical protein,” rather than a function demonstrated to be involved in horizontal gene transfer).

**Figure 1.**
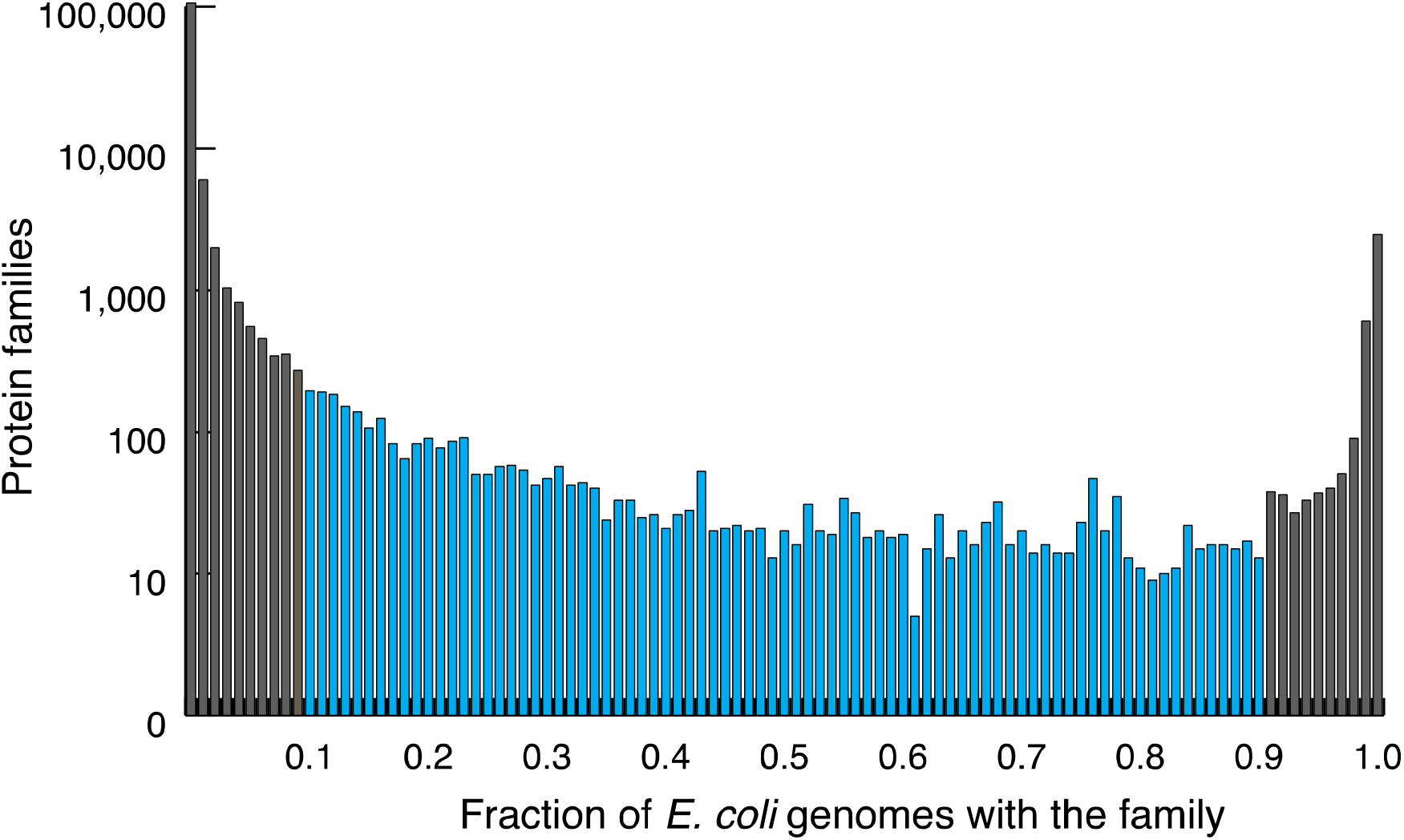
Histogram of the frequency of occurrence for all protein families found in the *E. coli* genomes in the BV-BRC. Models in this study were built to predict the presence or absence of protein families occurring in 10-90% of all genomes, shown in blue.

In order to reliably predict the presence or absence of each variable gene in each *E. coli* genome, a set of 100 highly conserved genes, present in all of the *E. coli* genomes (**Table S1**) was used to generate a feature set of nucleotide 7-mer counts. One XGBoost classifier was built for each of the 3,259 variable genes to predict its presence or absence. The models were trained and tested on a high-quality set of 4,000 *E. coli* genomes that was down sampled from all of the *E. coli* genomes in the BV-BRC. The training set includes 534 distinct MLSTs and 133 genomes that are untyped (53) (**Figure 2A, Table S2, Figure S1**). The F1 scores averaged across all 3,259 protein families was 0.912 ± 0.910-0.914 (± 95% confidence interval over 5 folds), with median F1 score of 0.926 (**Table S3**). When the F1 scores are averaged by genome or MLST, rather than by protein family, we observe similar results with F1 scores equal to 0.944 [0.943-0.945] per genome and 0.918 [0.913-0.923] per MLST (**Table 1, Figure 2C**).

**Table 1.** Macro F1 scores averaged by genome, MLST, and protein family.

| **Averaged by:** | **Training set**<br>**(4,000 genomes)** | **Hold-out set**<br>**(419 genomes)** |
| --- | --- | --- |
| Genome | 0.944 [0.943-0.945] | 0.880 [0.876-0.882] |
| MLST | 0.918 [0.913-0.923] | 0.867 [0.862-0.873] |
| Protein family | 0.912 [0.910-0.914] | 0.718 [0.712-0.724] |

**Figure 2.**
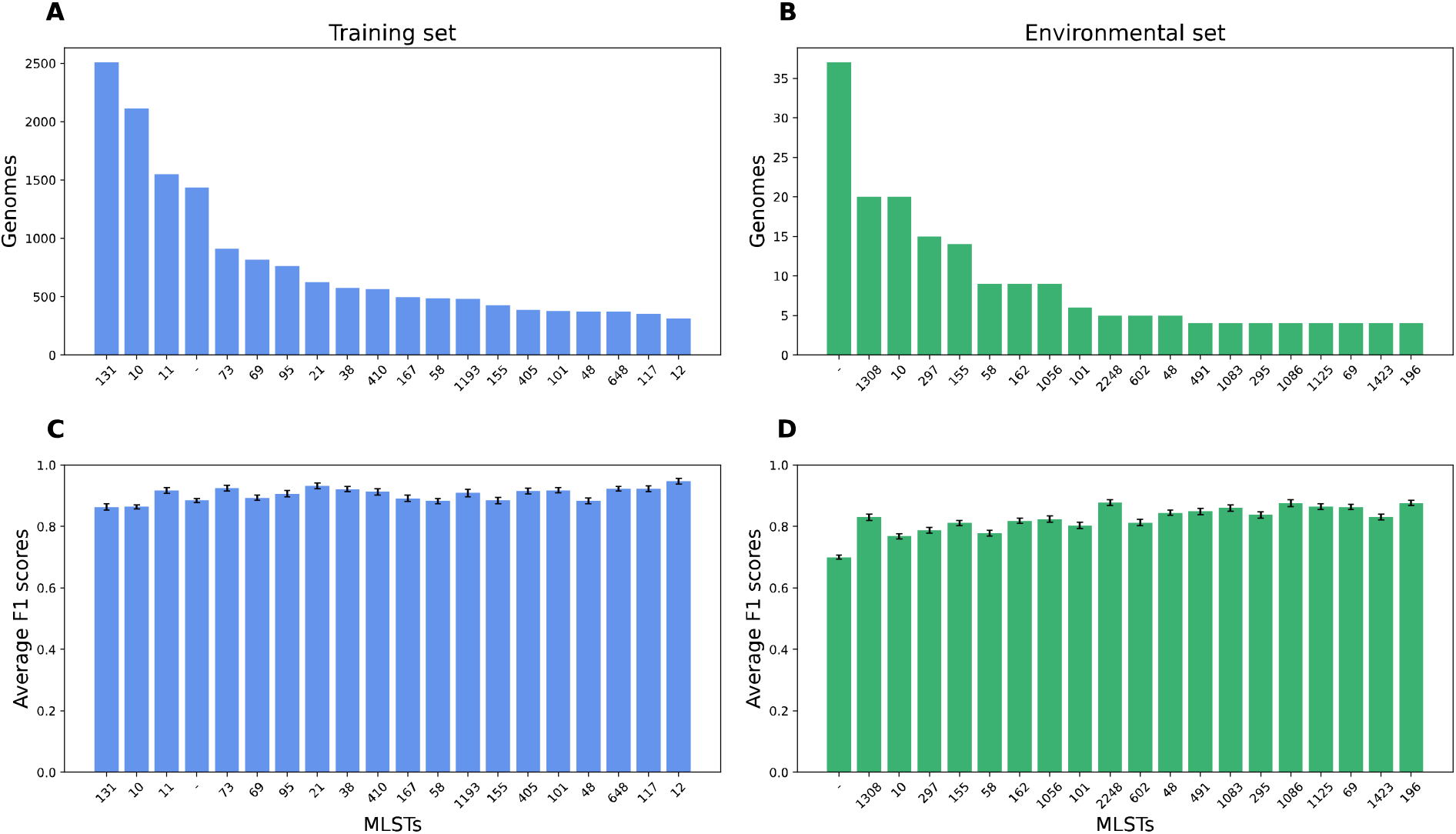
MLST distributions and F1 scores averaged by MLST. A) Histogram of the 20 most frequently occurring MLSTs in the training set of 4000 diverse genomes from the BV-BRC; B) Histogram of the 20 most frequently occurring MLSTs in the holdout set of 419 environmental genomes; C) F1 scores averaged by MLST for the set of 4,000 BV-BRC genomes; D) F1 scores averaged by MLST for the holdout set of environmental genomes. Error bars depict the 95% confidence intervals. The MLST labeled with a dash represent all genomes with undetermined MLSTs in each set.

Although the high F1 scores with cross validation indicate that the models are robust, models built for longer genes could have higher accuracies than shorter genes because the *ab initio* gene callers have difficulty accurately predicting shorter open reading frames (59). Likewise, genes that occur more frequently across the training set may have distribution patterns that are more consistent with the phylogeny of the conserved genes that were used as features, making their models more accurate. This might explain why the F1 scores are slightly higher when they are averaged by genome or MLST, because the more commonly occurring families are contributing more to these averages. To assess these potential sources of error, we plotted the average F1 scores for each protein family versus the median protein length for the protein family members and observe a weak correlation between gene length and accuracy PCC = 0.173 (**Figure 3A**). Similarly, when we plot F1 versus the occurrence of each family across the training set of 4,000 genomes, we observe a slightly upward trend in the average F1 scores with a PCC of 0.612. This trend is not dramatic, and the genes occurring least frequently, in 10-11% of the genomes, still have an average F1 score of 0.885 [0.866-0.904] (**Figure 3B**). Although these data indicate weak trends in model accuracy relating to protein length and abundance in the training set, this does not appear to be a major source of bias that could explain the high F1 scores that we observe.

**Figure 3.**
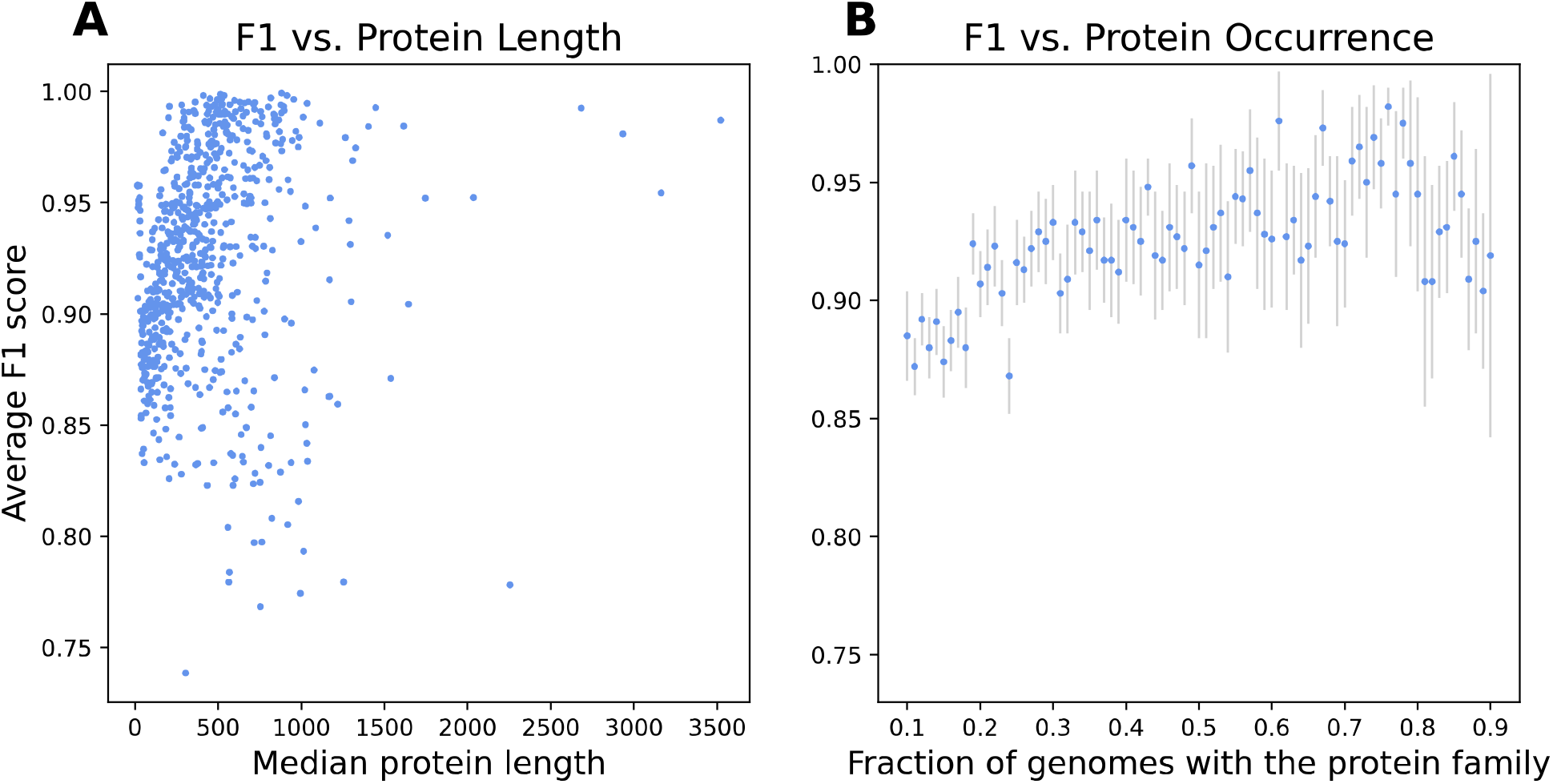
Average F1 scores versus protein length and protein family occurrence. A) F1 scores averaged by protein family plotted by the median protein length for all family members; B) F1 scores averaged by protein family versus the fraction of *E. coli* genomes in the training set containing a member of the given protein family. Gray bars depict the 95% confidence intervals.

### Models built with less data retain accuracy

In order to understand how using less data influences the performance of the models, we first built models using 7-mers from the top 25, 50, and 75 core genes. As expected, the models that were based on 25 core genes performed slightly worse (F1 = 0.886 ± 0.882-0.889, averaged by protein family) because they contain less information and gradually improved as the number of core genes was increased (**Figure 4A**). Likewise, we built models using the original set of 100 core genes as features, and gradually increased the size of the training set from 500 to 4,000 diverse *E. coli* genomes. The models trained on 500 genomes had an F1 score of 0.863 ± 0.859-0.867 (averaged by protein family), and the F1 scores gradually increased beyond 0.9 as the models were trained with 4,000 genomes (**Figure 4B**). This improvement is likely due to the better representation of the variable genes across the training set. Overall, the data suggest that reliable models can be built with fewer conserved genes or training set genomes with a correspondingly modest decrease in performance. Unless otherwise stated, results reported in this study are for models built from a feature set of 100 core genes and a training set of 4,000 *E. coli* genomes.

**Figure 4.**
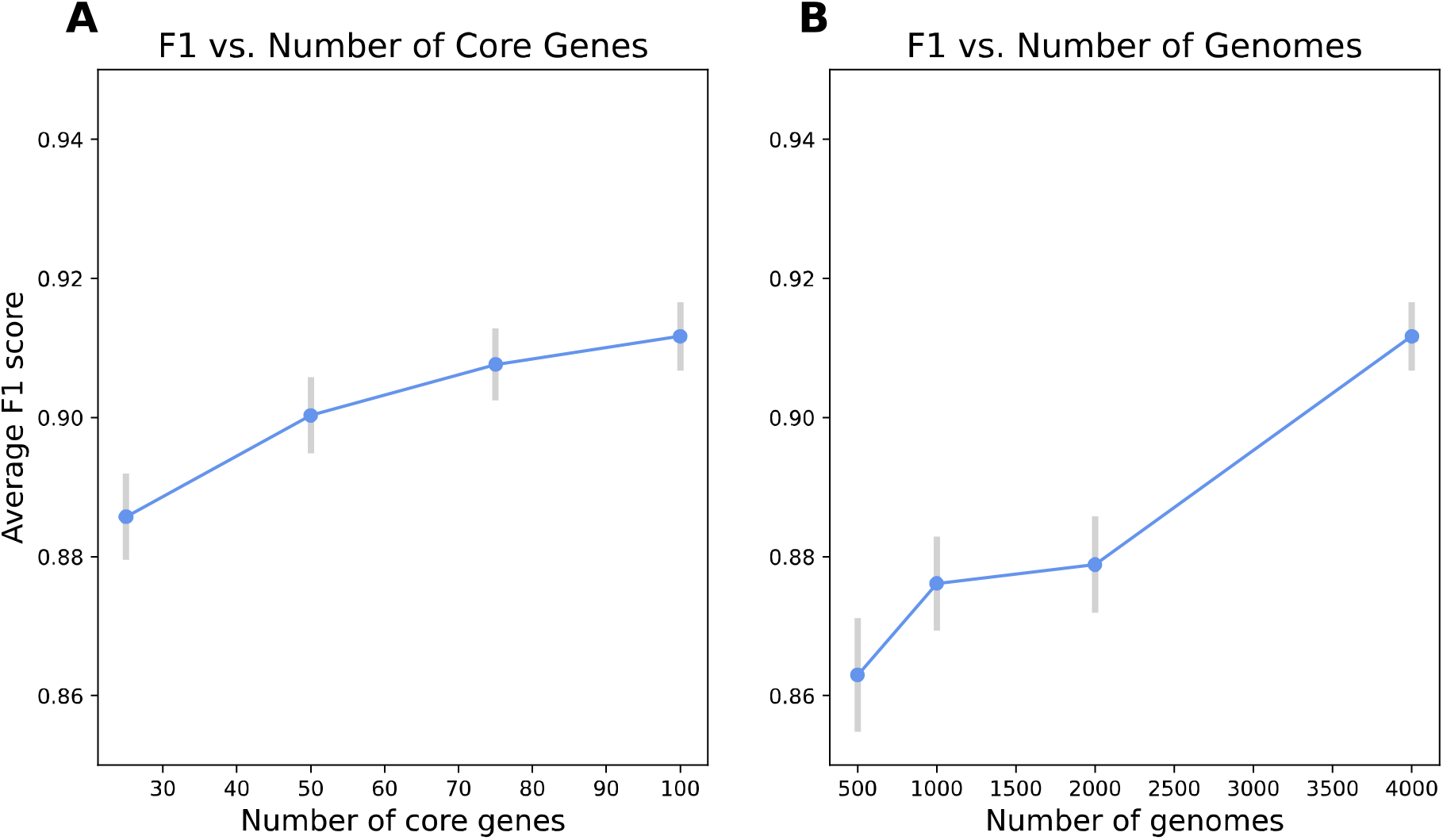
F1 scores versus number of core genes and number of genomes used to train the models. A) F1 scores averaged by protein family versus the number of core genes used to train the model; B) F1 scores averaged by protein family versus the number of diverse *E. coli* genomes used to train the models. Gray bars depict the 95% confidence intervals over 5 folds.

### Horizontally transferred genes can be predicted

Since the feature set for the models are based on conserved genes, it is possible that models for certain protein families outperform others due to their tight coupling to the phylogeny or may underperform due to the effects of horizontal gene transfer. When we examine the F1 scores based on the protein functions encoded by the variable genes, we find that the accuracy of the models is typically higher in genes that are well annotated (**Table 2, Table S3**). For instance, models for variable genes with functions occurring in subsystems (F1 = 0.935 ± 0.931-0.940), or which have full Enzyme Commission (EC) numbers (F1 = 0.945 ± 0.937-0.952) have significantly higher F1 scores than those that do not. Conversely, genes that are annotated with functions involved in horizontal gene transfer, including those encoding functions relating to transposable elements (F1 = 0.895 ± 0.882-0.907), phage elements (F1 = 0.872 ± 0.868-0.876), or conjugation and other plasmid-related functions (F1 = 0.824 ± 0.814-0.834) all had had significantly lower F1 scores than the genes that did not (**Table 2**). A total of 14 AMR-related protein families were modeled, and they have an average F1 score of 0.841 [0.814-0.869], indicating that they resemble the horizontally transferred genes (**Table S6**). These results indicate that although protein families with horizontal gene transfer-related functions do have lower F1 scores than other variable genes, their presence or absence can still be reliably predicted (F1 > 0.8).

**Table 2.** Commonly occurring protein functions in the set of 3,259 modeled protein families with their average F1 scores^1^.

| **Annotations** | **With the annotation** |  | **Without the annotation** |  |
| --- | --- | --- | --- | --- |
|  | **number** | **avg F1** | **number** | **avg F1** |
| Hypothetical etc. proteins<sup>2</sup> | 1343 | 0.902 [0.898-0.906] | 1916 | 0.918 [0.915-0.921] |
| Occurring in subsystems | 679 | 0.935 [0.931-0.940] | 2580 | 0.905 [0.903-0.908] |
| With the term "phage" | 513 | 0.872 [0.868-0.876] | 2746 | 0.919 [0.916-0.922] |
| With complete EC numbers | 245 | 0.945 [0.937-0.952] | 3014 | 0.909 [0.906-0.912] |
| Transporters | 166 | 0.949 [0.940-0.959] | 3093 | 0.910 [0.907-0.912] |
| Membrane proteins | 133 | 0.954 [0.945-0.962] | 3126 | 0.910 [0.907-0.912] |
| With the term "secretion" | 119 | 0.965 [0.959-0.971] | 3140 | 0.910 [0.907-0.912] |
| Transcriptional regulation<sup>2</sup> | 97 | 0.945 [0.934-0.956] | 3162 | 0.911 [0.908-0.913] |
| With the term "fimbriae" | 78 | 0.974 [0.964-0.984] | 3181 | 0.910 [0.908-0.913] |
| Toxins and anti-toxins | 70 | 0.932 [0.917-0.946] | 3189 | 0.911 [0.909-0.914] |
| With the terms "transposase" or "transposon" | 67 | 0.895 [0.882-0.907] | 3192 | 0.912 [0.910-0.915] |
| With the terms "conjugation" or "plasmid" | 59 | 0.824 [0.814-0.834] | 3200 | 0.913 [0.911-0.916] |
| Relating to flagellar function | 39 | 0.948 [0.943-0.952] | 3220 | 0.911 [0.909-0.914] |

<sup>1</sup>The average is reported with the 95% confidence interval
<sup>2</sup>Containing the terms "hypothetical", "uncharacterized," "putative," or "mobile element"
<sup>3</sup>Containing the terms, transcriptional "activator," "repressor," "regulator," or "antiterminator"

### Model performance on an environmental holdout set

Although the collection of public *E. coli* genomes is large, it is biased toward laboratory, surveillance, and clinical strains. We wanted to observe how well the models trained on the public genomes would extend to novel genomes. To do this, we sequenced a collection of 419 environmental *E. coli* isolates that were collected from freshwater environments. Importantly, none of these genomes previously existed in the public archives. Overall, the collection is comprised of 136 distinct MLSTs and 37 untyped genomes, and the distribution of MLSTs differs from that of the public collection (**Figure 2B, Figure S1**). When the models that were trained on the public data are applied to these genomes, we observe F1 scores of 0.880 [0.876-0.882] averaged by genome, 0.867 [0.862-0.873] averaged by MLST (**Figure 2D**), and 0.718 [0.712-0.724] averaged by protein family (**Table 1**). The diverse genomes lacking an MLST designation, have a lower average F1 score of 0.700 [0.693-0.706], and is likely due to their genetic diversity which has not been learned by the models. Likewise, the lower F1 scores averaged by protein family are likely due to the differences in distribution of the protein families across these genomes. For instance, approximately 38% of the protein families that existed in 10-90% of the public genomes occur in less than 10% of the genomes in the environmental collection (**Figure S2**). In other words, the distribution of these protein families deviates from the expetaction of the models trained on the public data. However, since these these families are rare within the envioronmental collection, they are insufficient to dramatically alter the results when averaged by genome or MLST (**Figure S3**), both of which remain greater than 0.86. Overall, these results indicate that the models are robust for predicting variable gene content in holdout set of diverse environmental *E. coli* genomes.

## Discussion

In this study, we predicted variable gene content across *E. coli* genomes by building machine learning classifiers to predict the presence or absence of each protein family occurring in 10-90% of the high-quality genomes for the species. In order to make the predictions, we used nucleotide k-mers from a set of core genes that were held in common by all members of the species. Overall, the average F1 scores were greater than 0.9 over the training set of 4,000 diverse genomes, indicating that the data from the conserved genes is sufficient for predicting the presence or absence of many of the variable genes. When we looked at how the accuracy relates to protein functions, we found that genes that were well annotated, either belonging to a SEED subsystem or annotated as having complete EC numbers were more easily predicted, since they had significantly higher F1 scores than those that did not. Conversely, models for genes with functions associated with horizontal gene transfer had significantly lower F1 scores. Although this is unsurprising given that horizontal gene transfer moves these genes in patterns that do not necessarily match the phylogeny, it is noteworthy that genes with annotations containing the terms “plasmid” and “conjugation,” which was the category with the lowest average F1 score in our analysis, still had a remarkably high average F1 score of 0.824. This is likely due to the ability of XGBoost to track non-linear relationships. Another surprise was that the models for genes with protein functions containing the terms “hypothetical,” “uncharacterized,” and “putative” had average F1 scores of 0.902 and were only slightly lower than the F1 scores for the set of families with curated annotations. This suggests that despite being poorly characterized, the occurrence of these sequences is rather easily predicted, implying that there is much more to learn about their distribution patterns and value to be added by elucidating their functions.

Using a holdout set of 419 *E.* coli genomes from freshwater environmental isolates, the models retained extensibility with F1 scores of 0.880 and 0.867 averaged by genome and MLST respectively, indicating that the models work well even in diverse genomes. In both the training set and the holdout set, we observed slight correlations in the accuracy of each model and the underlying protein length and occurrence of each family. However, these trends were insufficient to explain the high F1 scores for the models. The influence of rare families was more dramatic in the holdout set lowering the F1 score averaged by protein family to 0.718. However, since almost 40% of the families occurred in less than 10% of the genomes in the holdout set, their per-genome effect was considerably smaller. Adding diverse genomes to the training set as they become available would eventually correct this issue.

One limitation of this study is that by focusing on the set of variable genes occurring in 10-90% of the genomes, many of the rarely occurring genes were omitted. Although predicting the presence or absence of this massive and enigmatic set of genes is obviously desirable, this was done to control the study size and because these rare genes often lacked sufficient numbers to provide balanced sets for modeling. As long as the *E. coli* pan genome remains open and we continue to observe new genes with each new genome (41, 42), this will always be a problem, so predicting the presence or absence of the rarest genes may require a different modeling strategy. However, we expect that as the number of genomes increases, the number of protein families that can be used to build balanced classifiers in the way that we did in this study will also continue to increase.

Our highest quality set of models was generated using 100 core genes as features on a training set of 4,000 *E. coli* genomes and covered the set of protein families occurring in 10-90% of the genomes. This resulted in a collection of over 3,259 XGBoost models. This approach represented a rather significant outlay of computing resources, with each model taking approximately 4 minutes on an Intel Xeon Gold 6148 machine utilizing 128 cores, for a total of 7.6 days for computing the entire set. Although this experimental design is admittedly brute force, it is nevertheless tractable and could be extended to other well sequenced species. Indeed, unlike previous models that we have built for predicting AMR phenotypes using larger k-mer sizes and more complex matrices (36, 60, 61), these models are simple binary classifiers and have small memory footprints, and thus could be computed in parallel on a cluster with a modest amount of memory per node, rather than a high memory server. In designing this study, we attempted several other matrix designs and algorithms, including several deep learning approaches which had the potential to make the task more succinct. However, these attempts have been unsuccessful in our hands due to the size of the dataset, and ultimately the strategy of computing one classifier per family was successful. One way to reduce the computational burden might be to use fewer core genes or training set genomes. We showed that systematically reducing the size of the training set, while maintaining diversity, or using a smaller number of core genes for the feature set resulted in modest losses in accuracy. These tradeoffs may be deemed acceptable in certain circumstances. Using this study as a proof of concept, we expect that future studies will find more elegant modeling solutions.

In conclusion, we have found that it is possible to predict the presence or absence of a large number of the *E. coli* variable genes by building classifiers that use k-mers from a set of conserved genes. These models were highly accurate and worked even for families with hypothetical and unknown functions. This study provides a potential framework for predicting whether a genome or MAG should or should not be expected to contain a given gene, and has implications for the estimation of genome quality, the assessment of risk due to AMR and other virulence genes, and the ability to predict the presence of other important genes.

## Abbreviations

AMR: Antimicrobial Resistance
BV-BRC: Bacterial and Viral Bioinformatics Resource Center
MAG: Metagenome assembled genome
MLST: Multi Locus Sequence Type
PATRIC: PAThosystems Resource Integration Center
RAST: Rapid Annotation Subsystem Technology
XGBoost: Extreme Gradient Boosting

## Acknowledgements

We thank Emily Dietrich for her careful editing and Bob Olson for technical assistance. This work was funded in part by the United States National Institute of Allergy and Infectious Diseases Bacterial and Viral Bioinformatics resource center award [Contract No. 75N93019C00076] to PI Rick Stevens, and by the United States Defense Advanced Research Projects Agency iSENTRY Friend or Foe program award [Contract No. HR0011150042] to JJD, the National Science Foundation Awards [MCB-1616737 and DBI-1229585] to AAB, and the Herbert H. and Grace A. Dow Foundation. The funders had no role in study design, data collection and interpretation, or the decision to submit the work for publication.

