## Supplemental Figures for "Predicting Protein-encoding Gene Content in *Escherichia coli* Genomes"

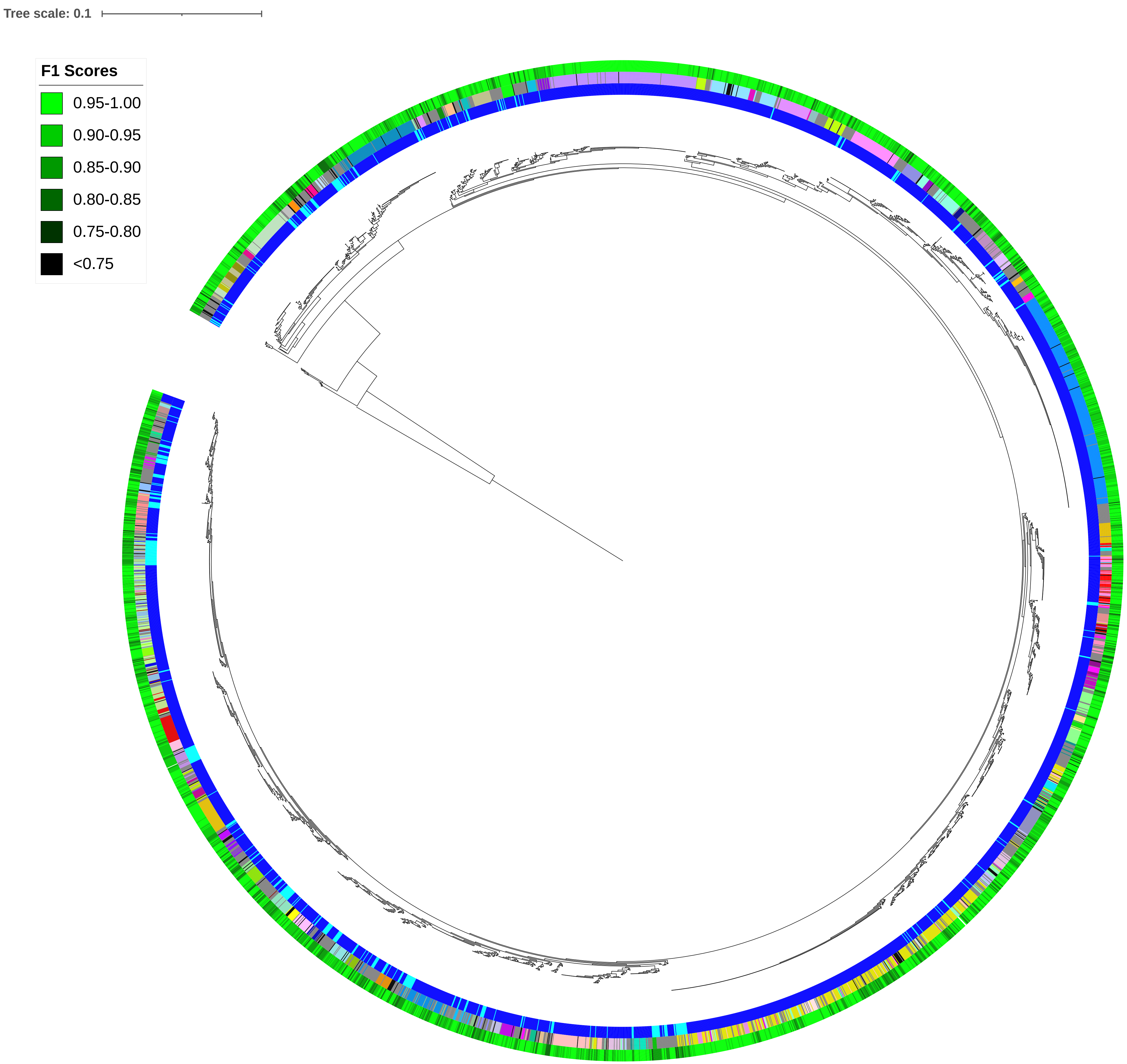

Tree scale: 0.1
F1 Scores
0.95-1.00
0.90-0.95
0.85-0.90
0.80-0.85
0.75-0.80
<0.75

**Figure S1.** Phylogenetic tree of *E. coli* strains used in this study. The tree was built from a concatenated nucleotide alignment of the protein-encoding genes from the 5 most conserved families in **Table S1**. The tree is rooted on *Salmonella enterica* Typhimurium LT2. The inside ring depicts the 4,000 *E. coli* genomes from the training set (blue) and the 419 environmental strains (teal). The middle ring depicts the 100 most common MLSTs as different colors, with the remainder shown in gray. The outside ring depicts the average macro F1 score for each genome with the color scale shown in the top left.

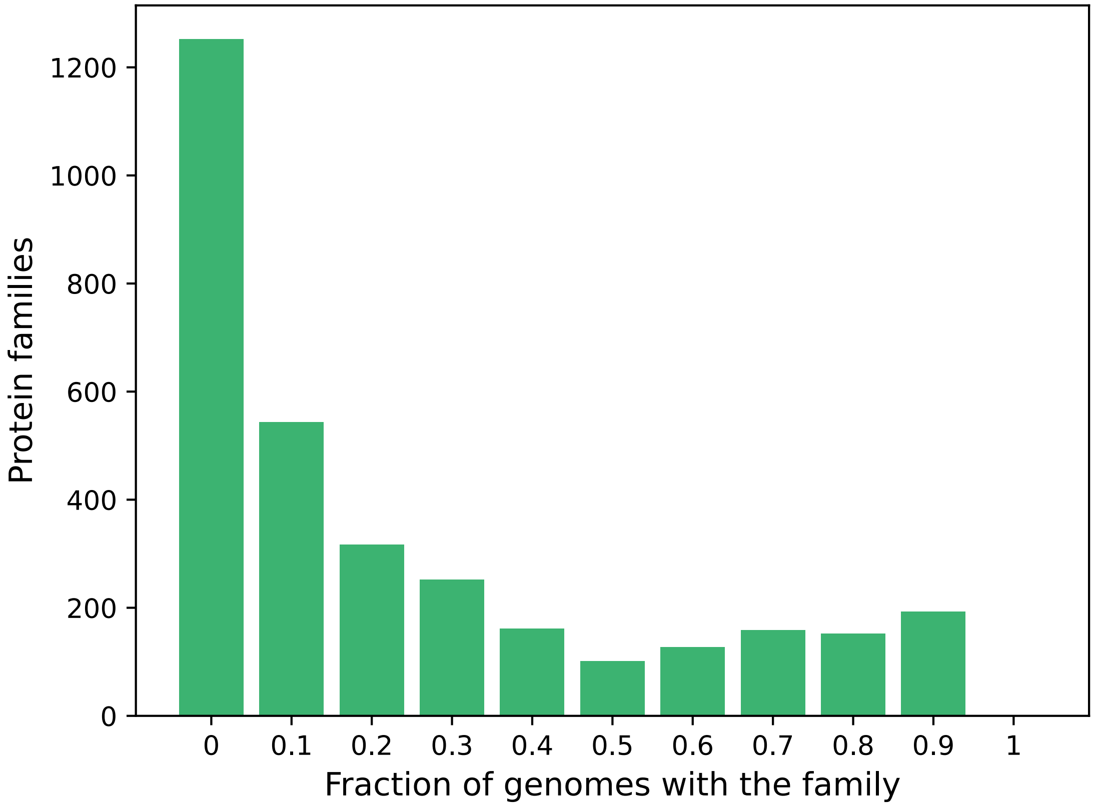


**Figure S2.** Histogram depicting the fraction of 419 environmental *E. coli* genomes containing each protein family that was modeled in this study.

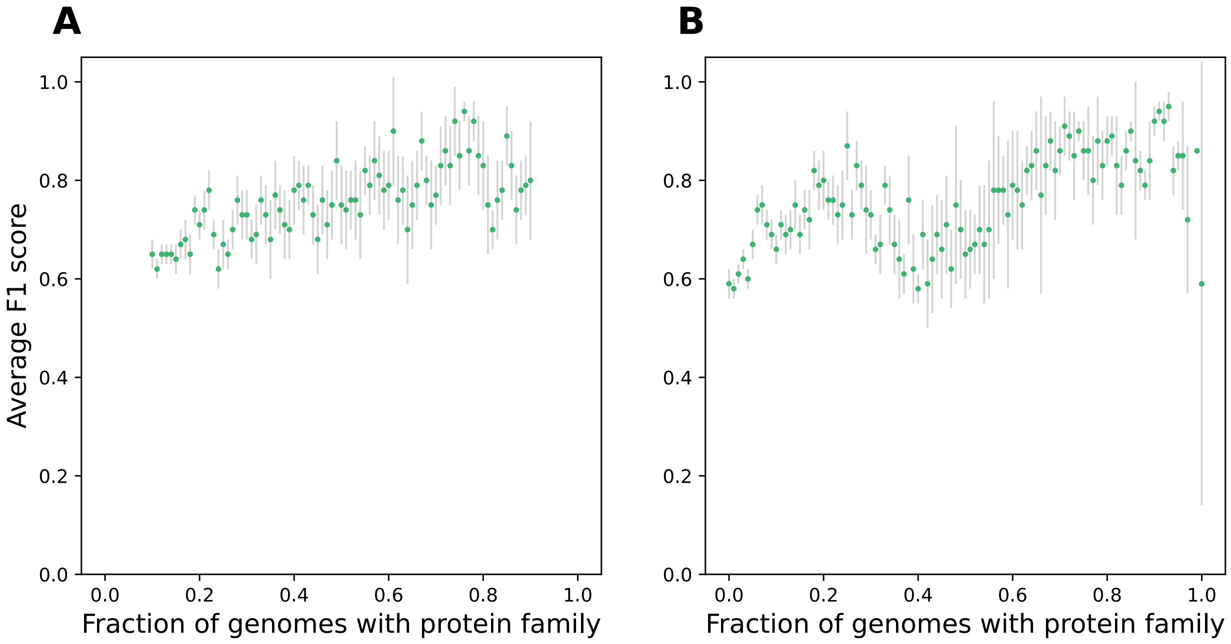


**Figure S3.** Average F1 scores for each protein family model in the environmental set versus the fraction of *E. coli* genomes containing a member of the given protein family. In A) the X-axis depicts the fraction of the training set genomes from the BV-BRC containing each protein family, and in B) the X-axis depicts the fraction of the 419 environmental *E. coli* genomes containing each protein family. Gray bars depict the 95% confidence intervals.
